# Oligomerisation mediated by the D2 domain of DTX3L is critical for DTX3L-PARP9 reading function of mono-ADP-ribosylated androgen receptor

**DOI:** 10.1101/2023.11.29.569193

**Authors:** Carlos Vela-Rodríguez, Chunsong Yang, Heli I. Alanen, Rebeka Eki, Tarek A. Abbas, Mirko M. Maksimainen, Tuomo Glumoff, Ramona Duman, Armin Wagner, Bryce M. Paschal, Lari Lehtiö

## Abstract

Deltex proteins are a family of E3 ubiquitin ligases that encode C-terminal RING and DTC domains that mediate interactions with E2 ubiquitin-conjugating enzymes and recognise ubiquitination substrates. DTX3L is unique among the Deltex proteins based on its N-terminal domain architecture. The N-terminal D1 and D2 domains of DTX3L mediate homo-oligomerisation, and the D3 domain interacts with PARP9, a protein that contains tandem macrodomains with ADP-ribose reader function. While DTX3L and PARP9 are known to heterodimerize, they assemble into a high molecular weight oligomeric complex, but the nature of the oligomeric structure, including whether this contributes to the ADP-ribose reader function is unknown. Here, we report a crystal structure of the DTX3L N-terminal D2 domain and show that it forms a tetramer with, conveniently, D2 symmetry. We identified two interfaces in the structure: a major, conserved interface with a surface of 973 Å^2^ and a smaller one of 415 Å^2^. Using native mass spectrometry, we observed molecular species that correspond to monomers, dimers and tetramers of the D2 domain. Reconstitution of DTX3L knockout cells with a D1-D2 deletion mutant showed the domain is dispensable for DTX3L-PARP9 heterodimer formation, but necessary to assemble an oligomeric complex with efficient reader function for ADP-ribosylated androgen receptor. Our results suggest that homo-oligomerisation of DTX3L is important for mono-ADP-ribosylation reading by the DTX3L-PARP9 complex and to a ligand-regulated transcription factor.

## Introduction

Ubiquitination is a post-translational modification mediated by the consecutive actions of three enzyme families (Hershko & Ciechanover, 1998; Scheffner et al., 1995). The modification controls a multitude of physiological and disease-associated pathways. This enzymatic cascade results in a covalent linkage between ubiquitin, a 76 amino acid protein, and a target protein (Hershko & Ciechanover, 1998). A protein can be modified on only one residue or simultaneously on multiple residues, processes referred to as mono-ubiquitination and multi-mono-ubiquitination, respectively (Yau & Rape, 2016). Ubiquitin itself can be modified on one of eight amino acid residues, thus generating a poly-ubiquitinated protein (Dittmar & Winklhofer, 2019; Komander & Rape, 2012; Tracz & Bialek, 2021; Yau & Rape, 2016). The length and the linkage type of the poly-ubiquitin chains determine the fate of the modified protein. For example, polyubiquitin chains linked through lysines 48, 63, or 11 can direct a protein for proteasomal degradation (Chowdhury et al., 2023; Ohtake et al., 2018; Pickart, 1997; Zheng et al., 2023; Zhu et al., 2023).

In addition to regulating protein half-life, ubiquitination can also modulate protein function by influencing localisation, activity and interactions with partner proteins (Magits & Sablina, 2022; Tracz & Bialek, 2021). Protein ubiquitination can be regulated through the catalytic function of de-ubiquitinating enzymes (DUBs) or through allosteric mechanisms. By hydrolysing the isopeptide bond, DUBs remove the mono-ubiquitin moiety or the poly-ubiquitin chains from a protein, thereby counteracting ubiquitination actions and restoring the pool of free ubiquitin (Clague et al., 2019; Schober & Berra, 2016; Snyder & Silva, 2021). Ubiquitination is also regulated through the coordinated action between the E2 conjugating enzymes and E3 ubiquitin ligases or through oligomerisation of ubiquitin E3 ligases (Kiss et al., 2021; Plechanovová et al., 2012). This is most widely observed in RING-type E3 ligases in which dimerization results in the activation of CHIP and TRIM E3 ligases (Das et al., 2022; Fiorentini et al., 2020; Koliopoulos et al., 2016; Nikolay et al., 2004) as well as in enhanced activity, such as the case of the BRCA1-BARD1, Mdm2-MdmX and DTX3L-DTX1 heterodimers (Brzovic et al., 2003; Kosztyu et al., 2019; Linke et al., 2008; Mallery et al., 2002; Stewart et al., 2018; Takeyama et al., 2003).

Deltex3-like (DTX3L) is one of the five human Deltex (DTX) proteins, which have a common domain at the C-terminus, responsible for substrate recognition and facilitating ubiquitin discharge from the E2 conjugating enzyme (Obiero et al., 2012; Takeyama et al., 2003). DTX1-4 share a middle proline-rich region, speculated to be involved in protein-protein interactions. DTX1, 2 and 4 also bear a tandem of N-terminal WWE domains, suspected to act as mono-ADP-ribose (MAR) and poly-ADP-ribose (PAR) reading modules (Aravind, 2001; DaRosa et al., 2015). DTX3L has a distinctive domain architecture as it substitutes the WWE domains and proline-rich region with three domains termed D1, D2 and D3 (**Figure 1A**). It has been established that in addition to DTX1, DTX3L interacts with the HECT-type E3 ubiquitin ligase AIP4 as well as the ADP-ribosyltransferases PARP9 and PARP14 (Vela-Rodríguez & Lehtiö, 2022).

**Figure 1.**
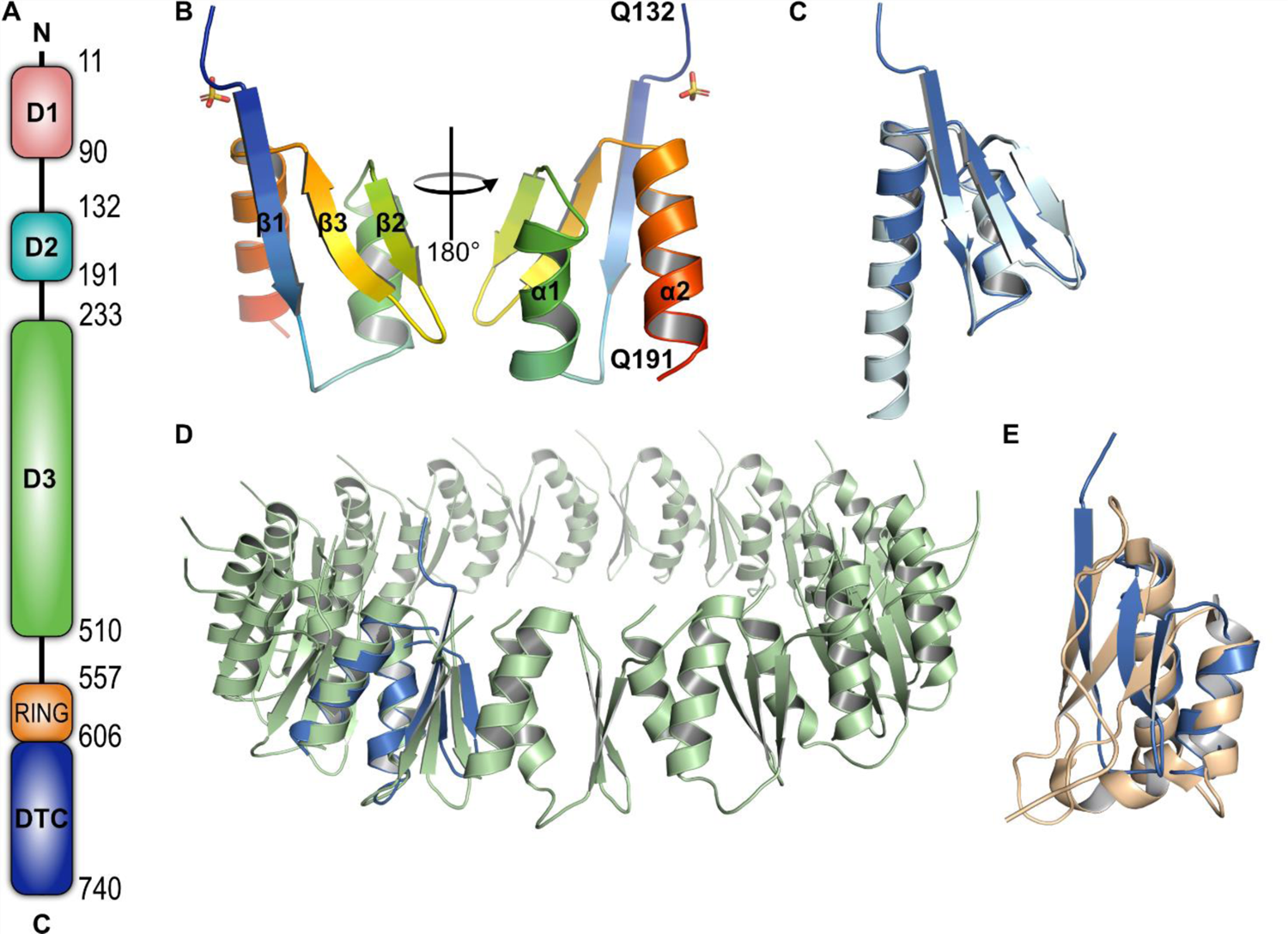
Solved structure of D2 domain of DTX3L is very similar to the predicted structure of AlphaFold2 and is present in other oligomerisation modules. A) Schematic of the domain architecture of DTX3L. B) Cartoon representation of the determined crystal structure of DTX3L D2 domain in rainbow colour. Secondary structure features are labelled according to the amino acid sequence. C) Superimposition of experimentally solved structure (marine) against the predicted structure by AlphaFold2 (cyan), RMSD = 0.853 Å, 56 Cα atoms. D) Superimposition of the experimentally determined structure of DTX3L D2 against residues 237-317 of the type II secretion system protein D (light green; PDB: 5ZDH), RMSD = 3.7 Å, 59 Cα atoms. E) Superimposition of the experimentally determined structure of DTX3L D2 against KH domain 1 of E3 ubiquitin ligase MEX3C (wheat; PDB: 5WWX), RMSD = 2.9 Å, 52 Cα atoms.

Our previous studies have shown that the DTXL3 D3 domain mediates the interaction with PARP9, protein that grants the complex the MAR-reading function present in DTX1, 2 and 4 (Ashok et al., 2022). We also established that DTX3L can form high-molecular weight oligomers mediated by its D2 domain and that an oligomeric-deficient construct had increased auto-ubiquitination activity *in vitro* (Ashok et al., 2022). PARP7-mediated MARylation of the AR and binding to DTX3L-PARP9 contributes to androgen-dependent gene regulation (Yang et al., 2021). A study on the MAR-reading function of PARP9 on an androgen receptor-(AR) derived peptide indicates that this function is significantly enhanced for the oligomeric DTX3L-PARP9 complex compared to isolated PARP9 or an oligomerisation-deficient DTX3L-PARP9 complex (Wijngaarden et al., 2023). Based on this information, the oligomerisation of DTX3L is not only a structural feature but also plays an active role in its functionality.

In the present study, we describe a crystal structure of the isolated D2 domain of DTX3L. We show that D2 assembles as a homo-tetramer formed by a dimer of dimers associated in a head-to-tail orientation. To gain a better understanding of the significance that D2-mediated oligomerisation has in cells, we analysed the impact of this feature of the DTX3L-PARP9 complexes in prostate cancer cells that express the AR (PC3-AR) (Yang et al., 2021). The experiments in which the cells were treated with synthetic androgen to promote MARylation of the AR correlate with previous studies establishing that the oligomerisation of the DTX3L-PARP9 complex is required for the recognition of MARylated AR.

## Results

### D2 domain shares structural features to type II secretion protein D and KH domains

We initially aimed to crystallise the N-terminal region of DTX3L (amino acids 1-200) based on our previously defined boundaries (Ashok et al., 2022) **Figure 1A**. After several crystallisation efforts, we subjected the protein to *in situ* proteolysis with chymotrypsin. The treatment facilitated the formation of polyhedral crystals of ca. 120 µm in length. Using mass-spectrometry, we identified that the amino acid sequence of the crystallisable fragment started at A126 and ended in Q200. Preliminary data collection at the home source diffractometer revealed that the crystals were highly anisotropic, which remained even after optimisation of the crystallisation conditions. Taking advantage of the presence of methionine and cysteine residues in the protein sequence we solved the structure by sulphur single-wavelength anomalous diffraction (S-SAD) using high-redundancy data collected on the long-wavelength beamline I23, at Diamond Light Source, UK (**Table 1**).

**Table 1.** Data collection and refinement statistics.

| PDB ID: 8R79 |  |
| --- | --- |
| **Data collection** |  |
| Resolution ($\text{\AA}$) | 70.446-2.181 (2.344-2.181) |
| Wavelength ($\text{\AA}$) | 2.7552 |
| Beamline | DLS, I23 |
| Temperature (K) | 80 |
| Space group | P 4<sub>1</sub>2<sub>1</sub>2 |
| Cell dimensions |  |
| *a*, *b*, *c* ($\text{\AA}$) | 140.893 140.893 41.037 |
| $\alpha$, $\beta$, $\gamma$ ($^\circ$) | 90, 90, 90 |
| No. of unique reflections | 12554 (628) |
| *R*<sub>merge</sub> | 0.107 (3.117) |
| Mean *I*/σ*I* | 33.3 (1.7) |
| Spherical completeness (%) | 56.5 (15.0) |
| Ellipsoidal completeness (%) | 91.7 (67.2) |
| Redundancy | 90.6 (55.2) |
| CC<sub>1/2</sub> | 1.000 (0.842) |
| **Refinement** |  |
| *R*<sub>work</sub> / *R*<sub>free</sub> | 0.210 / 0.249 |
| No. non-H atoms | 1948 |
| protein | 1928 |
| ligands/ions | 20 |
| waters | 0 |
| *B*-factors ($\text{\AA}^2$) | |
| protein | 58.33 |
| ligands/ions | 64.16 |
| waters | 0 |
| RMSD bonds / angles | 0.007 / 1.51 |
| Ramachandran plot (%) |  |
| favored/allowed/outliers | 99.14 / 0.86 / 0.0 |

Despite the anisotropy, we observe a well-defined electron density for the protein (**Figure S1**), for which we determined that the D2 domain comprises three antiparallel β-sheets packed against two α-helices (**Figure 1B**). Notably, the experimental structure aligns with residues 136 to 191 of the AlphaFold2 prediction with an RSMD of 0.853 Å (**Figure 1C and Figure S2**). While the alignment between the helices and the β2 and β3 strands is almost perfect, α2 and β1 are 9 amino acids shorter and 3 amino acids longer, respectively, in the experimentally determined structure. Noteworthy, the AlphaFold2 server was released after we solved the structure, and in retrospect, it could have served as a model in molecular replacement.

To identify if the fold was present in other proteins, we used the Dali server to look for structural similarities in the protein data bank (PDB) (Holm et al., 2008). The search provided over 600 results with a Z score ranging from 2.0 to 5.9 and sequence identity to the identified protein structures lower than 20% (**Table S1**). The top-ranking characterised protein was the type II secretion protein D (PDB: 5ZDH). The chain to which the structure was aligned was part of an oligomeric assembly. While both structures shared the triple strands and double helix arrangement, it was interesting to notice that, unlike our solved structure, all the strands of the secretion protein were 5 amino acids long (**Figure 1D**). Furthermore, this strand-shortening feature was similar to KH domains 1 and 2 of the E3 ubiquitin ligase MEX3C (PDB: 5WWW and 5WWX, respectively) (**Figure 1E**). Dali identified both KH domains with a score of 3.8 and a sequence identity of 11 and 10%, respectively.

### D2 domain of DTX3L assembles in a homo-tetramer

The asymmetric unit of the crystal had four copies of the D2 monomer, and the interfaces buried in this oligomer (973 Å^2^ and 415 Å^2^) indicate that this could be the biologically relevant oligomer and that D2 assembles as a homo-tetramer with a 2-fold symmetry between adjacent dimers (Figure 2A). Notably, we identified a large positive density in the centre of the symmetry axis, which we could not explain as a component of the crystallisation solution. The surface representation suggests that the tetramer forms a tight cavity (Figure 2B). Despite that the maximum resolution of the data was 2.18 Å, crystal anisotropy and low high-resolution completeness compromised the quality, and we did not model any water molecules. We, however, identified four sulphate ions derived from the ammonium sulphate of the crystallisation conditions. Sulphates were located at the N-terminus of helix α2 and each of them form a hydrogen bond with H185 (Figure 1B **& 2A,C**).

**Figure 2.**
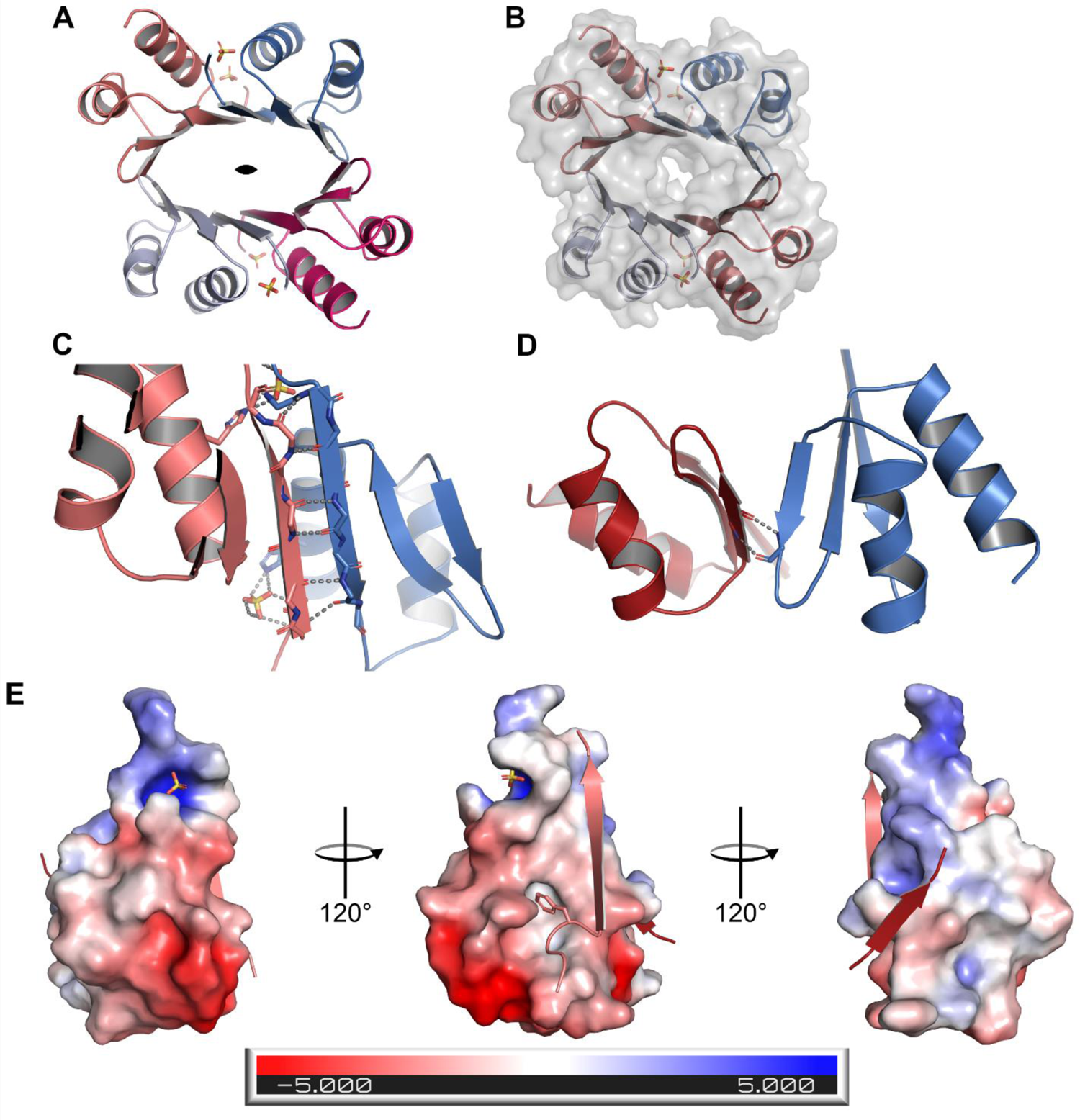
D2 domain of DTX3L assembles as a homo-tetramer with 12 antiparallel β-strands forming a barrel. A) Upper view of the D2 homo-tetramer displaying the barrel-like conformation. Chains are represented in different colours and the two-fold symmetry symbol marks the symmetry axis. B) Surface of the tetramer shows a tight interface C) Main interaction interface is arranged by complementary antiparallel β helices. Grey dashes indicate the polar contacts between chains and between the chain and its conjugated sulphate ion. D) Secondary interaction interface between symmetrical dimers is formed by weak antiparallel strands. Grey dashes indicate the hydrogen bonds. E) Electrostatic potential surface for a monomer with its complementary β1 and β2 strands. In the β1 strand complementary to the ESP and F135 are shown as sticks.

The larger monomer-monomer interface corresponds to the head-to-head arrangement between the β1 strands of adjacent monomers. This assembly continues the antiparallel β sheet of the monomer with six hydrogen bonds between the strands (Figure 2C). The smaller interface is formed between the short β2 strands and only two inter-chain hydrogen bonds are formed (Figure 2D). The electrostatic surface of the monomer shows that the larger interface is not only formed by the hydrogen bonds of the antiparallel strands, but that the interaction surface is mostly hydrophobic (Figure 2E**, left**). Conversely, PISA analysis shows that the smaller interface has a combination of hydrophobic and polar interactions between chains (Figure 2E**, right**). At the main interface there is a hydrophobic cleft that accommodates Ile134 and Phe135 from its complementary chain (Figure 2E**, left**). Taking all into account, dimer and tetramer are stable in solution, but the structural features suggest that the dimer might be a more stable assembly. This correlates with the free energy of assembly dissociation (ΔG^diss^), calculated to be 0.8 kcal/M for the tetramer and 19 kcal/M for the dimer.

### Native mass spectrometry (MS) and small-angle X-ray scattering (SAXS) confirm that D2 is an oligomer in solution

Crystallisation experiments were performed at a high protein concentration, which could generate artefactual oligomers. To determine if the D2 oligomeric behaviour was preserved in solution, we conducted native MS experiments with the protein. By conducting the experiment in denaturing conditions, i.e. diluting the protein in 50% acetonitrile and formic acid instead of ammonium acetate, the calculated mass for the protein was 11.3 kDa, correlating with the theoretical molecular weight (11.5 kDa). On the other hand, experiments that were conducted in native conditions showed a mixture of monomer, dimer and tetramer based on the molecular weight (Figure 3A). To further confirm that D2 is an oligomer in solution, we analysed the sample with size exclusion coupled SAXS. From the normalised Kratky plot, we observe a single peak whose maximum almost crosses the intersection of the axes, a behaviour characteristic of proteins that are mostly globular (Figure 3B). In the same line, the Porod exponent was calculated to 3.7. The SAXS analysis did not only reveal that D2 was mostly globular, but that the molecular weight also corresponded to that of a tetramer with 43 and 47 kDa from ScÅtter (Classen et al., 2013) and SAXSMow (Piiadov et al., 2019), respectively.

**Figure 3.**
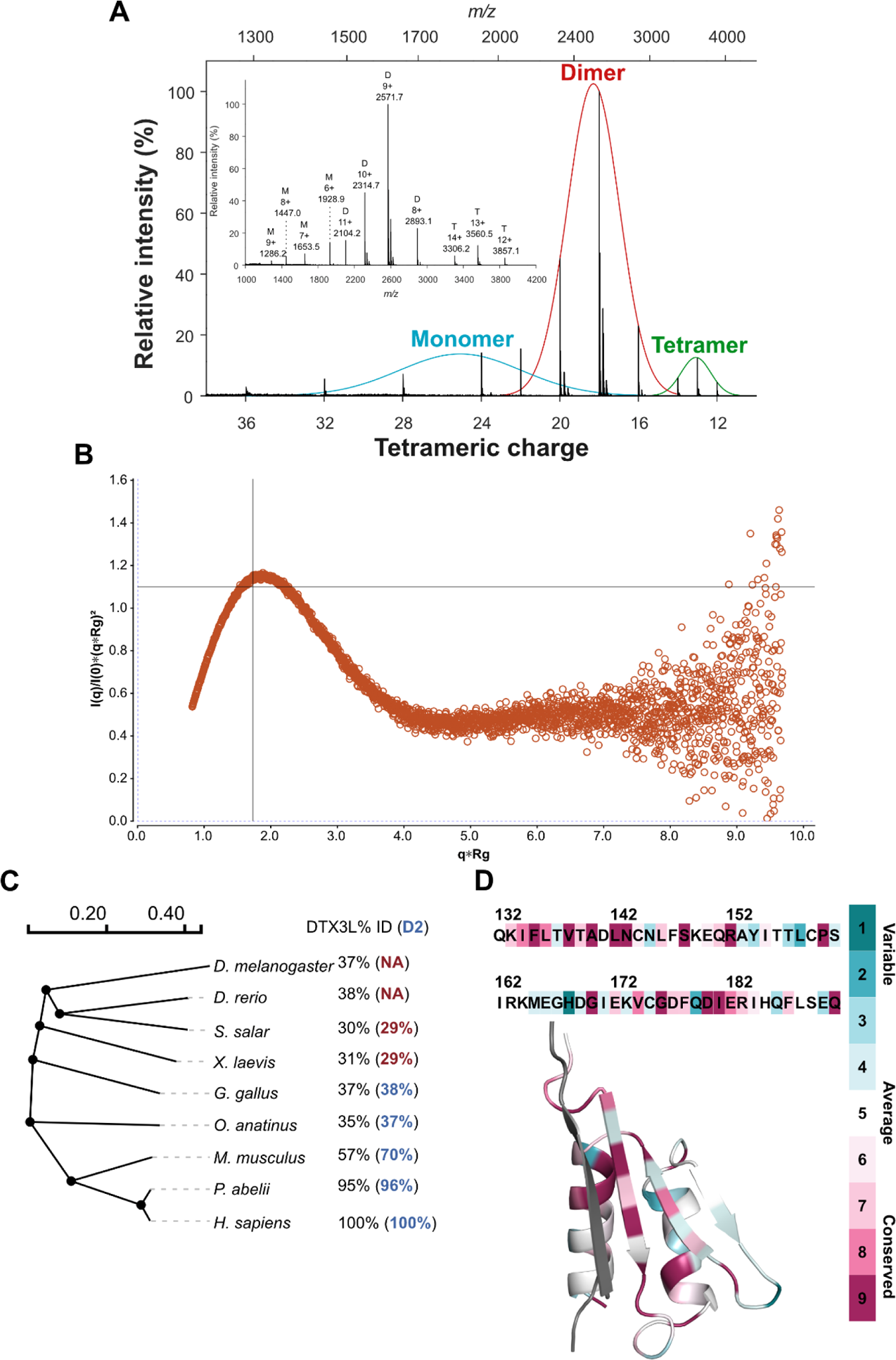
D2 oligomer is also observed in solution and is the main interaction interface is conserved throughout multiple species. A) Native mass spectrometry deconvoluted spectra of DTX3L D2 identifies monomeric, dimeric and tetrameric species. B) Dimensionless Kratky plot of DTX3L D2 (I101-Q200). C) Slanted dendogram constructed of the global sequence alignment of DTX3L from model organisms that are representative of different taxonomy levels. The percentage of identity of each organism to human DTX3L is annotated on their right in black. The per cent of sequence identity of a global pairwise sequence alignment between human DTX3L D2 and each respective organism is show in blue, for organisms with identity lower than 30% the value is shown in red. D) Estimated evolutionary conservation of DTX3L D2 using chain A of the solved structure as a query in Consurf. The amino acids of the chain are coloured according to the conservation scale shown in the right and the complementary strand of the dimer is coloured in grey.

### The principal dimer interface of D2 is a highly conserved characteristic of DTX3L in amniotes

We were not only interested in seeing the behaviour of DTX3L D2 in solution, but also in seeing how this feature was conserved from an evolutionary perspective. We, therefore, first identified the sequence identity between human DTX3L full-length amino acid sequence and its orthologues in model organisms, which represent different taxonomy levels (Figure 3C). From the sequence identity, we were able to conclude that DTX3L, similar to PARP9, is not a particularly conserved protein (Sowa et al., 2022). Furthermore, the identity between orthologues was mostly observed in the RD regions (Figure 1A) of the protein while the remaining domains showed a variable degree of identity. A variable N-terminus is also noticeable by the lack of a D2-like region in *D. melanogaster* and *D. rerio* and a low degree per cent identity in *S. salar* and *X. laevis*.

An interesting feature that we observed was that in *S. salar* and *X. laevis* a sequence resembling the second half of the D2 domain, corresponding to I612-Q192 of the human orthologue, was observed but not anything that could be paired with the N-terminal region of the domain. It is not until the emergence of amniotes, that this sequence can also be identified, which reflects in a higher identity. Moreover, by analysing the protein sequence with the ConSurf server (Ashkenazy et al., 2016), we noticed that the main interaction interface of the dimer had a higher conservation score than the rest of the domain (Figure 3D). This is reflected mainly in the amino acids that form the hydrophobic core of the interface, including Phe135, indicating that this residue could be a key point in the assembly of the dimer.

### D2-mediated oligomerisation of DTX3L and PARP9 promotes recognition of ADP-ribosylation

The native DTX3L-PARP9 complex from prostate cancer cells, as well as when the complex is reconstituted from recombinant proteins, is a multimer that by gel filtration is predicted to contain 5-6 copies of the DTX3L-PARP9 heterodimer (Ashok et al., 2022; Yang et al., 2017). In crosslinking experiments, D1 has a tendency to form dimers and D2 forms tetramers (Ashok et al., 2022). PARP9 contains two MAR binding macrodomains and the DTX3L-PARP9 complex would therefore be endowed with at least eight macrodomains. This arrangement is interesting since AR MARylation by PARP7 occurs on multiple Cys sites, including seven sites within the unstructured N-terminal domain (NTD) of AR (Yang et al., 2021). Given these considerations, we posited that AR-DTX3L-PARP9 assembly might reflect multi-valent engagement of PARP9 macrodomains with ADP-ribosyl-Cys groups within AR. To test this model, we assessed the reader activity of PARP9 co-expressed with DTX3L^WT^, and with mutant DTX3L^ΔN^ (DTX3LΔ2-229). The mutant heterodimerises with PARP9 but fails to undergo oligomerisation (Ashok et al., 2022). We prepared DTX3L Knockout (DTX3L^KO^) cells in the prostate line PC3-AR, reconstituted the cells with DTX3L^WT^ or with DTX3L^ΔN^, and examined complex formation with MARylated AR by immunoprecipitation and immunoblotting to detect PARP9 binding (reader function). Cells stably reconstituted with WT DTX3L show a robust level of endogenous PARP9 binding to ADP-ribosylated AR, which depends on PARP7 since the interaction is lost when cells are treated with the PARP7 inhibitor RBN2397 (Figure 4A**, lanes 1, 2**). By contrast, only low levels of PARP9 binding to ADP-ribosylated AR are observed in cells that lack DTX3L (+Vector), or in cells reconstituted with the DTX3L mutant deficient for oligomerization (+DTX3LΔN). Notably, PARP9 associates with both WT DTX3L and DTX3L^ΔN^, showing that the N-terminal domain of DTX3L is dispensable for PARP9 binding (Figure 4B). Overall, the data argue that oligomerization is important for efficient binding of DTX3L-PARP9 to MARylated-AR. As an additional test of the model, we prepared MARylated AR and used it for binding assays with recombinant DTX3L-PARP9 oligomers and DTX3L^ΔN^-PARP9 heterodimers (Figure 4C). Similar to the IP results with DTX3L KO cells, recombinant PARP9 displayed only a low level of binding to MARylated AR, and this was augmented by preassembly with full-length recombinant DTX3L (Figure 4D). From these data we conclude that oligomer formation mediated by the DTX3L D1-D2 region increases the binding efficiency of PARP9 macrodomains that read multi-site MARylation.

**Figure 4.**
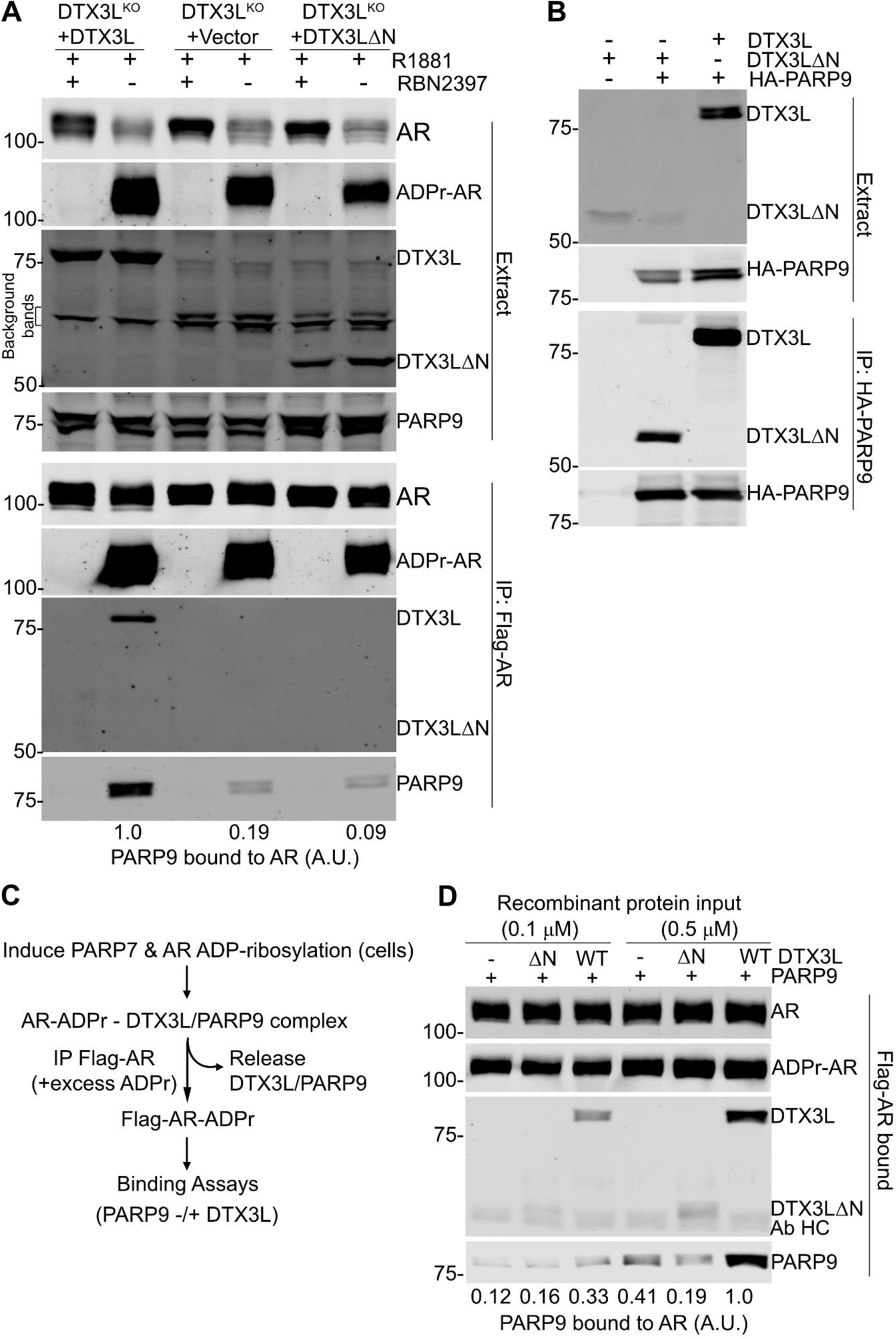
Multimerization of DTX3L-PARP9 is critical for reading AR ADP-ribosylation. A) Reconstitution of MAR reading on AR in DTX3L^KO^ cells. PC3-AR-DTX3L^KO^ cells were stably transduced with WT DTX3L, vector alone, and DTX3LΔN. Cells were treated with synthetic androgen (R1881) to induce PARP7 and AR MARylation; the PARP7-selective inhibitor RBN2397 was included as a control to prevent AR MARylation. Flag-AR complexes were isolated by immunoprecipitation and probed for MAR (Kamata et al., 2019; Yang et al., 2021), and for DTX3L and PARP9 as described (Yang et al., 2017). Bound PARP9 (%WT) was normalized with MAR-AR. The level of PARP9 binding to AR (arbitrary units, A.U.) are shown relative to binding with full-length DTX3L. B) Multimerization mutant DTX3L^ΔN^ forms a complex with PARP9. PC3-AR-DTX3L^KO^ cells were transiently transfected with the indicated constructs, and binding to PARP9 was detected by immunoprecipitation and immunoblotting. The DTX3L^ΔN^ is only recovered in the immunoprecipitated products when cells are co-transfected with HA-PARP9. C) Scheme for in vitro AR/DTX3L-PARP9 complex reconstitution. D) Reconstitution of MAR-reading on AR in vitro. Flag-AR from PC3-AR (HA-PARP7) cells treated with R1881 (2 nM, 6 h) was immunoprecipitated on magnetic M2-beads in the presence of 0.5 mM ADPr to deplete bound DTX3L-PARP9 from cells. Beads-immobilized ADP-ribosylated AR (MAR-AR) was used for the in vitro binding assays with recombinant PARP9 alone, PARP9-DTX3LΔN, and PARP9-DTX3L (recombinant protein inputs are shown in **Supplementary Figure S3**). The level of PARP9 binding to AR (arbitrary units, A.U.) are shown relative to the level of binding with full-length DTX3L (0.5 μM protein inputs). The position of antibody heavy chain (Ab HC) in the AR IP is indicated on the panel.

## Discussion

Our work is the first one to provide insights not only into the structural assembly of DTX3L, but also on the functionality that this multimer has at cellular level. The crystal structure revealed a tetrameric assembly of DTX3L D2 domain, which was only possible to crystallise after treating a longer construct with chymotrypsin. Taking the AlphaFold model into account (**Supplementary Figure S1**), there is a disordered region of nearly 40 amino acids connecting D1 and D2, which would explain why crystallisation trials for this construct were not successful. A stable oligomer would also explain why the crystallised fragment corresponded to D2 instead of D1. Our structure determination revealed two interaction interfaces between monomers, a larger interface driven through β1-β1’ and a secondary interface of β2-β2’ strands. It is in the main interface where we notice more deviations between the predicted and experimental structure, in which the latter one has a longer β1 strand and a shorter α2 helix. These features are complementary to each other as they form the groove in which Phe135 is positioned, and the longer strand at the N-terminus allows the formation of 2 additional hydrogen bonds which increase the stability of the dimer. According to the AF2 model, the second helix should contain also the last eight residues of the domain, but our data did not show electron density for them which suggests that they are instead disordered. The head-to-head assembly hints that the N- and C-termini of complementary chains could be facing each other.

While we could not identify the 12-stranded barrel assembly in the databases, not even when combining the chains in only one chain, the Dali server identified regions of different proteins that showed a similar fold as DTX3L D2 (**Table S1**). The results showed proteins with various functionalities, ranging from transport to enzyme activity. The KH domains of the E3 ubiquitin ligase MEX3C were among the most interesting results as it has been recently pointed out that they are found in several proteins from the PARP family, PARP9 included (Suskiewicz et al., 2023). In addition to the D2 domain, the D3 domain of DTX3L is predicted to be composed of four small domains resembling a KH fold. KH domains have been characterised as nucleic acid-binding modules present in transcription regulation, but also found to act as scaffolds for oligomerisation and protein-protein interactions (Valverde et al., 2008). The fact that the interaction between DTX3L and PARP9 is mediated by what has also been recently established as KH domains indicates that the KH domains in DTX3L are involved in protein-protein interactions (Ashok et al., 2022), including homo-oligomerization, instead of nucleic acid recognition.

The crystal structure of the D2 domain also reveals that the domain assembles as a tetramer with two distinct interaction interfaces, and the PISA analysis of the structure shows that while the tetramer of chains joined by the larger interfaces are energetically more stable, the tetramer is also a relevant biological unit. This would explain why in SAXS we could calculate the molecular weight of a tetramer but in native MS, we saw mainly dimeric species, as the energy required for the detection was enough to disrupt the β2-β2’ interface.

The fact that D2 emerged with amniotes and is particularly conserved in mammals suggest the idea that during evolution, DTX3L might have gained an oligomerising feature. We looked at the conservation of PARP14, PARP9, PARP7 to try to identify if DTX3L oligomerisation was related to the emergence of any of these proteins, which have been reported to form part of the interactome of DTX3L. We could not, however, establish a direct connection between the appearance of D2 and the gain of these proteins. Regarding the conservation of the amino acid sequence of the domain, we were surprised to see that while the percent of identity drops to 37 in birds, there are few amino acids that seem to be fully conserved throughout species. These amino acids were mainly located in the β1-β1’ interface and were mainly hydrophobic, which supports our hypothesis that the is interface is a well-characterised feature of DTX3L. Among these amino acids Phe135 is also amongst those with the highest conservation score assigned by Consurf. Considering the number of sequences that were considered for the analysis, it is possible to assume that the conservation of these residues is significant.

A recent study about the synthesis of androgen receptor-derived peptides with dual ADP-ribosylation sites showed that a complex of full-length DTX3L and full-length PARP9 had higher affinity towards the peptide than a complex in the D1-D2 domains of DTX3L were deleted (Wijngaarden et al., 2023). Our data correlates with the study by showing that PC3-AR cells expressing DTX3L^WT^ and PARP9 were bound to immunoprecipitated MARylated AR. On the other hand, levels of PARP9 bound to AR in cells that expressed oligomer-deficient DTX3L were significantly lower, demonstrating that the two macrodomains of PARP9 are not enough and the MAR-binding activity of PARP9 is dependent on oligomerisation of DTX3L. The enhancement of the binding has been attributed to the oligomer providing multiple copies of PARP9 and potentially their accessibility in the full-length context is limiting binding to the MARylated peptide in such a way that only independent copies of PARP9 are able to recognise modified residues that are close to each other.

## Materials and Methods

### Cloning

Coding sequence for DTX3L D2 (amino acids 101-200) was amplified by PCR and inserted by SLIC to a pNIC-MBP backbone and subsequently was transformed in *E. coli* NEB5α.

### Protein production

DTX3L D1-D2 and D2 were expressed in *E. coli* and purified as previously described (Ashok et al., 2022). For PARP9 expression in cells, we used the plasmid pKH3/HA-PARP9 (Yang et al., 2021).

### Crystallisation and data collection

DTX3L D1-D2 (1-200) was crystallised at 22°C in presence of chymotrypsin (1:100 mg/mL ratio) using a hanging drop vapour diffusion strategy. For crystallisation, 2 µL of protein (8 mg/mL; 380 µM) were mixed with 2 µL of precipitant solution [2.2 M (NH_4_)_2_SO_4_, 100 mM BisTris (pH 6.3)]. Star-like crystals appeared after 19 h and they were cryo-protected by adding 0.7 µL of 30% glycerol, 1 M (NH_4_)_2_SO_4_ on top of the crystal-containing drop. Immediately after, crystals were fished and flash-frozen in liquid nitrogen.

Data was collected on the long-wavelength beamline I23 at Diamond Light Source, at a wavelength of 2.755 Å, as five sweeps of 360 degrees, using the kappa goniometry to change the orientation of the crystal for each sweep. Integration and scaling were performed with autoproc (Vonrhein et al., 2011), using staraniso for anisotropy correction (Tickle et al., 2018). Experimental phasing by S-SAD was performed using the pipeline Crank2 (Skubák & Pannu, 2013), which yielded a partial model of the protein that was subsequently used as a template for molecular replacement with PHASER (Read & McCoy, 2011). Coot (Emsley & Cowtan, 2004) and REFMAC5 (Murshudov et al., 2011) were used for model building and refinement, respectively. The images of the structures were prepared using PyMOL (The PyMOL Molecular Graphics System, version 1.8.4.0, Schrödinger, LLC.).

### Identification of crystallisable fragment through Mass-spectrometry (MS)

Identification of the fragment was done through peptide mapping and denatured mass measurement. For peptide mapping, crystals were fished and washed with fresh crystallisation solution. They were subsequently washed with 2 µL of water and transferred to a micro-centrifuge tube. Sample volume was made up to 10 µL and subjected to SDS-PAGE. Band prepared and analysed with MALDI-TOF.

For denatured mass measurements, 2 µL of HPLC-grade water was added to drops containing crystals. The crystals were fished and transferred individually to 1.5 µL HPLC-grade water drops and thereafter to 1.5 µL drops of 10 mM ammonium acetate. Drops were aspirated with a pipette and the spot was rinsed with 1.5 µL of 10 mM ammonium acetate. Sample volumes were adjusted to 50 µL with 50% ammonium acetate and 50% acetonitrile. Mass was determined with Lumos.

### Native MS measurements

Protein was diluted to 10 mg/mL in 20 mM HEPES (pH 7.5), 150 mM NaCl, 0.5 mM TCEP and subsequently, buffer was exchanged to 200 mM ammonium acetate (pH 4.5) with a Zeba™ 7 K MWCO desalting column (ThermoFisher). After buffer exchange, the protein was diluted to 1 mg/mL in 200 mM ammonium acetate (pH4.5). Before the measurement, the diluted protein was filtered with a Vivaclear mini clarifying filter (0.8 µm PES) for 30 s at 6000 rpm. 9 µL from the sample were transferred to a NanoEs spray capillary (Cat no ES380; ThermoScientific). Measurements collected and analysed with BioPharma Finder (ThermoScientific).

### Small angle X-ray scattering data collection and analysis

D2 (10 mg/mL) were analyzed by SEC SAXS at Diamond Light Source B21 beamline (Oxfordshire, United Kingdom). Samples were run by injecting 50 µL of protein solution to a Superose 6 increase 3.2/300 column at a flowrate of 0.6 mL/min with 30 mM Hepes (pH 7.5), 350 mM NaCl, 5% glycerol, 0.5 mM TCEP. The column was connected to the SAXS system on the B21 beamline. Based on the Guinier Rg, matching frames from the main peak were averaged. Data processing was done using ScÅtter (Classen et al., 2013) to determine the maximum distance (Dmax), volume and Rg. Molecular weight was also calculated with the intensity file with the SAXSMoW server (Piiadov et al., 2019).

### Identification of similar structures in the PDB

The PDB of our structure was used as a query in the Dali server searching for similar structures in the PDB25 database as well as within the whole PDB (Holm et al., 2008). Due to the heuristic nature of the algorithm, the search was done in triplicates.

### Sequence identity and construction of phylogenetic tree

The amino acid sequence of human DTX3L (UniProt ID: Q8TDB6) was submitted as a query on Protein Path Tracker (Mier et al., 2018) to identify the earliest possible taxonomical family in which the protein was found. We retrieved the protein sequences of non-human orthologues by using human DTX3L as a query in the blastp suite (Hu & Kurgan, 2019) against the NCBI non redundant database. The search was performed using the organisms acquired from protein path tracker, predicted sequences were not taken considered for further analysis. We used Clustal Omega (Sievers & Higgins, 2014) to perform multiple sequence alignment of the retrieved sequences using the default parameters, the results were used to construct a phylogenetic tree with the neighbour-joining cluster method using the simple phylogeny tool (https://www.ebi.ac.uk/Tools/phylogeny/simple_phylogeny/ (accessed on May 29^th^ 2023)) from the EMBL-EBI web services. The presented tree was visualised with PRESTO (http://www.atgc-montpellier.fr/presto/<u>#</u> (accessed on May 29^th^ 2023)).

### Position-specific sequence conservation analysis

To identify the conservation degree for each amino acid of the crystallised protein, the refined structure was uploaded to the ConSurf server (Ashkenazy et al., 2016) using chain A as the base for the conservation analysis. Homologues were search with the HMMER algorithm with a minimal identity of 35% and an E-value of 0.0001. A total of 568 sequences were identified as HMMER hits, from which 268 were defined as unique and 150 (the maximum number of sequences allowed by the server) were used for the alignment. It was determined that JTT was the best evolutionary model for the data provided and conservation scores were calculated using the Bayesian method.

### Generation of PC3-AR cells with DTX3L deletion

To generate PC3-AR cells with DTX3L deletion, a pair of synthetic guide RNAs (sgRNAs; sg-DTX3L-1: 5’-CGGACTTGTACACCCGCACG-3’ and sg-DTX3L-2: 5’-CGGACTTGTACACCCGCACG-3’) was designed targeting distinct regions within Exon 1 of the *DTX3L* gene, encoding the DTX3L protein, using Benchling, and cloned into the Cas9-expresssing pX330 expression vector (Addgene #42230). Subconfluent PC3-AR cells were transiently transfected with the two Cas9-expressing plasmids (sg-DTX3L-1 and sg-DTX3L-2; 2.0 µg each), along with 0.5 µg pMSCVpuro vector (Clontech) containing a puromycin resistance gene, using Lipofectamine 2000 transfection reagent (Invitrogen); 24 hours following transfection, the cells were treated with puromycin (2 µg/ml) for 48 hours, after which single clones were isolated via serial dilutions. To identify clones with DTX3L deletion, DNA was extracted from individual clones for genotyping. Briefly, individual clones were grown in culture, and lysed overnight at 55°C in lysis buffer [100 mM NaCl, 10 mM Tris-HCl (pH 8), 25 mM EDTA, 0.5% SDS] supplemented with 20 µg of proteinase K. DNA was subsequently isolated using phenol chloroform/isoamyl alcohol extraction. Genotyping was performed via PCR amplification of the targeted DTX3L locus with primers flanking the two predicted Cas9 cleavage sites followed by Sanger sequencing (Eurofins). The following primers were used to amplify a 125 bp sequence spanning the two sgRNA target sites: DTX3L-F: 5’-AGAGCCATGGCCTCCCAC-3’, DTX3L-R: 5’-CCGCCCTCTCCCCTAGGTCA-3’. Immunoblotting of the individual DTX3L clonal knockout cells using an anti-DTX3L antibody was further used to confirm DTX3L deletion.

### Stable cell line generation using lentiviruses

Lentiviral vector pLH3 was constructed by replacing the U6 promoter in pLKO.1 GFP shRNA (Addgene #30323) with the CMV promoter in pKH3 (Addgene #12555) and additional cloning sites. DTX3L wild type (WT) and DTX3LΔN (DTX3LΔ2-229) were then cloned into the pLH3 vector. HEK293T (ATCC CRL-3216) were grown to ∼70% confluency and co-transfected with target plasmid (pLH3, pLH3/DTX3L or pLH3/DTX3LΔN) and accessory plasmids pMD2g and psPAX2 (ratio 2:1:1) using ViaFect transfection reagent (Promega E498A). After cell incubation at 37°C for ∼16 h, cells were replenished with fresh DMEM supplemented with 35% FBS, and further incubated for another 24 h. The medium containing lentiviruses was transferred to a fresh conical tube, clarified by centrifugation ∼700 x g, and then passed through a sterile 0.45 µm filter, with optional concentrating lentivirus using Takara’s Lenti-X Concentrator (Takara 631231). Infection of PC3-AR-DTX3L^KO^ cells were carried out in the growth medium (RPMI 1640 + 5% FBS + 1% Penicillin-Streptomycin solution) in the presence of 8 µg/ml of hexadimethrine bromide (Sigma H9268). After 2-3 cell doublings, the cells were selected with 1-2 µg/ml of puromycin.

### Immunoprecipitation

All steps were performed at 4°C. Cell pellets of PC3-AR with R1881 (2 nM, 17 h) and RBN-2397 (100 nM, 17 h) treatments were re-suspended in the cell extraction buffer [20 mM Tris-HCl (pH 7.5), 100 mM NaCl, 0.5% Triton X-100, 1 mM PMSF, 2 mM DTT, 5 mM EDTA, 5 µg/ml each of aprotinin/leupeptin/pepstatin with 0.5 µM Veliparib], and then end-over-end rotated for 20 min. The extracts were clarified with centrifugation (16,800 x g) for 20 min, and then subjected to anti-Flag M2 magnetic beads (Sigma M8823-5ML) or anti-HA magnetic beads (Pierce 88837) binding for 3-4 h. The beads were collected by magnetic fields, washed five times with the wash buffer [20 mM Tris-HCl (pH 7.5), 100 mM NaCl, 0.1% Triton X-100, 2 mM DTT, 0.1 mM EDTA, 1 µg/ml each of aprotinin/leupeptin/pepstatin, 0.5 µM Veliparib), and then re-suspended in SDS-loading buffer followed with Western blot analyses. Immunoblots were detected on an Oddysey CLx instrument (LI-COR) using the following reagents: anti-HA (HA.11 Clone 16B12 Monoclonal antibody, BioLegend 901514, used at 1:1,000), Alexa Fluor 680 donkey anti-rabbit IgG(H+L) (Invitrogen A10043) and goat anti-mouse IgG(H&L) DyLight 800 conjugated (Rockland 618-145-002).

### In vitro AR binding assays

To assemble DTX3L/PARP9 complexes, purified recombinant PARP9 was pre-incubated with DTX3L or DTX3LΔN (10 µM for each protein) for 6 h on ice. Other procedures were conducted at 4°C. Flag-AR was immunoprecipitated onto magnetic M2-beads from R1881 (2 nM, 6 h)-treated PC3-AR (HA-PARP7) cells in the cell extraction buffer supplemented with 0.5 mM ADPr to deplete bound DTX3L-PARP9 from the cells, and the beads were rinsed twice with the extraction buffer + 10 mg/ml BSA, washed 3 times (1 h each) with the extraction buffer + 10 mg/ml BSA + 0.5 mM ADPr, followed with five more washings (1 min each) with the extraction buffer + 10 mg/ml BSA (to deplete the free ADPr). Beads-immobilized ADP-ribosylated AR was then used for the overnight binding with recombinant proteins (PARP9 alone, PARP9/DTX3LΔN or PARP9/DTX3L; 0.1 µM or 0.5 µM in the extraction buffer + 10 mg/ml BSA), followed with 5 times washing with the extraction buffer. AR complex was released from the beads by heating at 95°C for 5 min in the SDS-loading buffer, and then subjected to SDS-PAGE and Western blot analysis.

## Data availability

Atomic coordinates and structure factors will be available at the Protein Data Bank with the id. 8R79. Raw diffraction data will be available at Etsin (https://doi.org/10.23729/b116653f-ddb5-4070-925e-c6e3260a8682). SAXS data is deposited to SASBDB.

## Author contributions

C.V.-R., B.M. P., L.L. conceptualization; C.V.-R., C.Y., H.I.A., R.E., T.A.A., M.M.M., R.D., A.W., investigation; C.V.-R., L.L. writing-original draft; C.V.-R, T.G., R.D., A.W., B.M.P., L.L. writing-reviewing and editing; B.M.P., L.L. funding acquisition; T.A.A., B.M.P, L.L. supervision.

## Acknowledgements

Biocenter Oulu Structural Biology core facility, member of Biocenter Finland, Instruct-ERIC Centre Finland and FINStruct, as well as of “Proteomics and Protein Analysis” and Sequencing core facilities are gratefully acknowledged.

