## Supplementary figures and tables for "Oligomerisation mediated by the D2 domain of DTX3L is critical for DTX3L-PARP9 reading function of mono-ADP-ribosylated androgen receptor"

### Contents

**Figure S1.** Representative electron density map.

**Figure S2.** Superimposition of D2 structure with AlphaFold model of DTX3L.

**Figure S3.** Recombinant protein inputs related to Figure 4.

**Table S1.** Results from a DALI search.

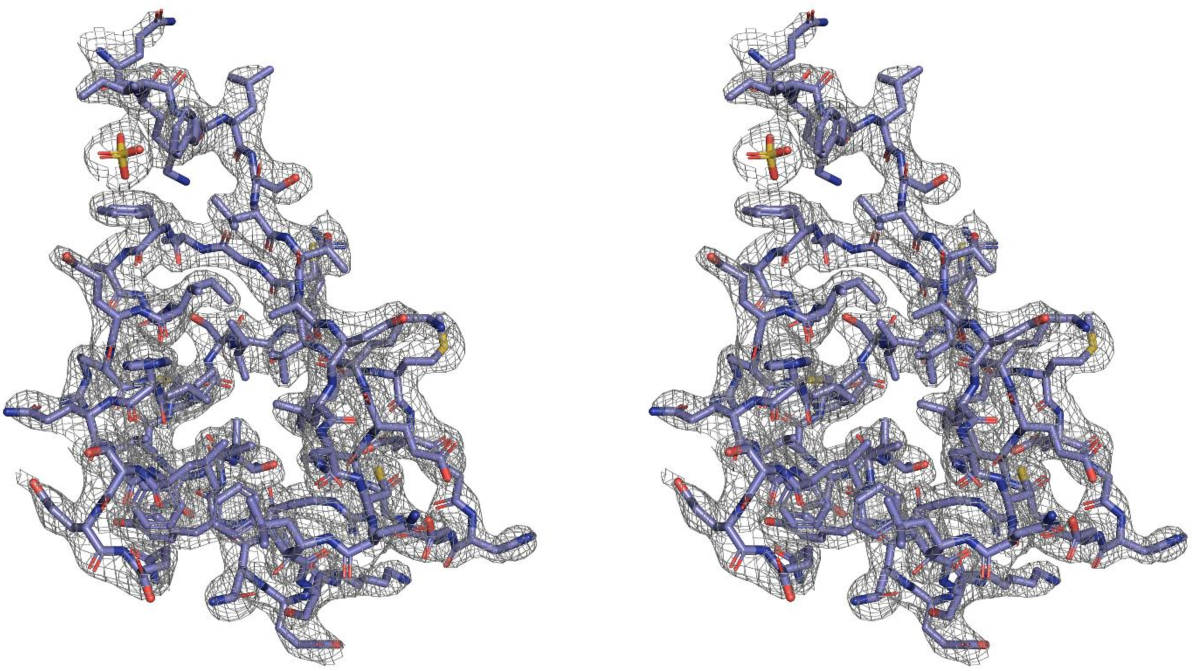

**Figure S1.** Stereoscopic representation of the electron density map of a monomer of the D2 domain of DTX3L. The electron density map is contoured at 1.0 sigma.

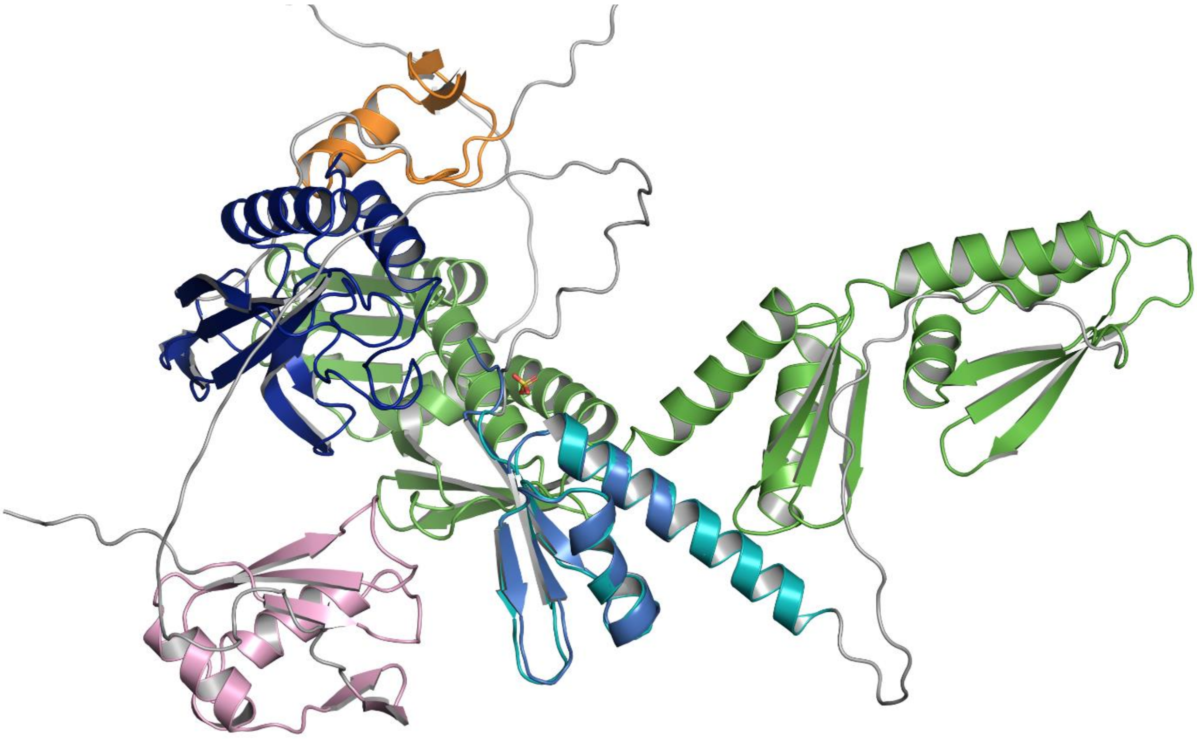

**Figure S2.** Superimposition of a monomer of the experimentally solved structure (marine) to the predicted full-length structure of DTX3L. Domains of DTX3L are colour-coded as the schematic in **Figure 1A**. AlphaFold2 prediction shows long, flexible regions between the domains of the protein.

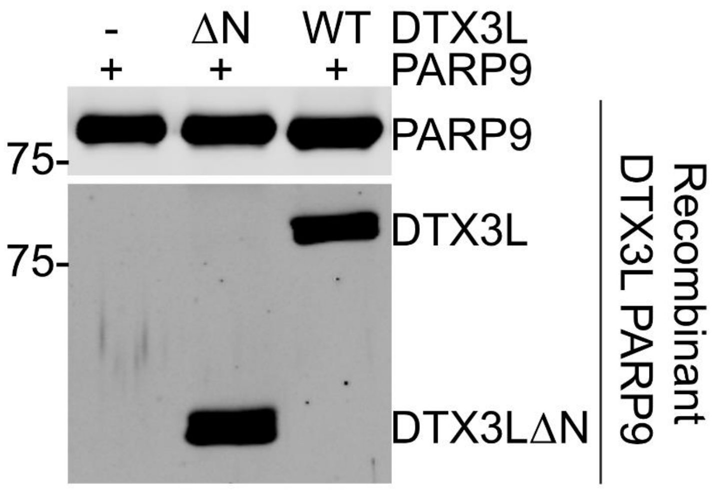


**Figure S3.** Recombinant protein inputs related to Fig. 4. Recombinant PARP9 alone or combined with DTX3L or DTX3LΔN (10 µM for each protein) was pre-incubated on ice for 6 h, and then diluted to 1 µM each with an extraction buffer [20 mM Tris-HCl (pH 7.5), 100 mM NaCl, 0.5% Triton X-100, 1 mM PMSF, 2 mM DTT, 5 mM EDTA, 5 µg/mL each of aprotinin/leupeptin/pepstatin with veliparib]. Equal volume of protein solution was further diluted with 1x SDS loading buffer and subjected to SDS-PAGE and Western blot analysis.

**Table S1.** Results obtained from the DALI against the PDB25 dataset.

| Chain | Z score | RMSD | Aligned residues | % id | PDB Description |
|---|---|---|---|---|---|
| 5zdh-A | 5.6 | 3.7 | 57 | 2 | TYPE II SECRETION SYSTEM PROTEIN D |
| 7eqx-A | 5.4 | 2.7 | 56 | 11 | CARBOXYPEPTIDASE B |
| 4ec5-A | 5.3 | 3.6 | 58 | 14 | GENERAL SECRETION PATHWAY PROTEIN D |
| 5hvf-A | 5.1 | 2.6 | 54 | 9 | CARBOXYPEPTIDASE B2 |
| 6w6m-A | 5.1 | 3.0 | 55 | 9 | TYPE IV PILUS SECRETIN PILQ FAMILY PROTEIN |
| 4av2-A | 5.1 | 2.5 | 58 | 9 | TYPE IV PILUS BIOGENESIS AND COMPETENCE PROTEIN P |
| 6n10-A | 5.0 | 3.4 | 57 | 9 | DIPHOSPHOMEVALONATE DECARBOXYLASE MVD1. PEROXISOM |
| 8h6s-A | 4.9 | 3.1 | 58 | 9 | MALONYL-COA-[ACYL-CARRIER-PROTEIN] TRANSACYLASE |
| 6u45-A | 4.9 | 3.3 | 55 | 11 | ELONGATION FACTOR 2 |
| 6wrv-C | 4.9 | 2.1 | 50 | 8 | TRANSFERRIN RECEPTOR PROTEIN 1 |
| 6wi5-A | 4.9 | 1.9 | 51 | 12 | DE NOVO DESIGNED PROTEIN FOLDIT4 |
| 6wrw-C | 4.9 | 2.2 | 49 | 8 | TRANSFERRIN RECEPTOR PROTEIN 1 |
| 3ce8-A | 4.8 | 2.4 | 52 | 8 | PUTATIVE PII-LIKE NITROGEN REGULATORY PROTEIN |
| 1jqg-A | 4.7 | 2.6 | 52 | 12 | CARBOXYPEPTIDASE A |
| 6fij-A | 4.7 | 3.1 | 54 | 9 | POLYKETIDE SYNTHASE |
| 2n2u-A | 4.6 | 2.1 | 51 | 12 | OR358 |
| 4qbu-A | 4.6 | 3.0 | 52 | 17 | ZMAA |
| 1i7q-A | 4.6 | 2.6 | 51 | 8 | ANTHRANILATE SYNTHASE |
| 5oqj-2 | 4.6 | 3.5 | 54 | 4 | DNA-DIRECTED RNA POLYMERASE II SUBUNIT RPB1 |
| 1pyt-A | 4.6 | 2.8 | 55 | 13 | PROCARBOXYPEPTIDASE A |
| 1in0-A | 4.5 | 2.5 | 50 | 10 | YAJQ PROTEIN |
| 6qum-N | 4.4 | 3.4 | 56 | 7 | V-TYPE ATP SYNTHASE ALPHA CHAIN |
| 4gx2-B | 4.3 | 2.9 | 53 | 9 | TRKA DOMAIN PROTEIN |
| 3cj8-A | 4.3 | 2.4 | 49 | 16 | 2.3.4.5-TETRAHYDROPYRIDINE-2.6-DICARBOXYLATE N- |
| 5d4o-A | 4.3 | 2.9 | 53 | 4 | NITROGEN REGULATORY PROTEIN P-II |
| 7lmx-A | 4.3 | 2.5 | 55 | 9 | INTEGRIN INHIBITOR |
| 7cyf-D | 4.3 | 2.8 | 52 | 8 | SLR1512 PROTEIN |
| 2pfd-A | 4.3 | 2.8 | 55 | 7 | FORMIMIDOYLTRANSFERASE-CYCLODEAMINASE |
| 3f56-B | 4.2 | 2.3 | 51 | 4 | CSOS1D |
| 5wx8-A | 4.2 | 2.8 | 56 | 2 | IMMEDIATE-EARLY PROTEIN 2 |
| 1wey-A | 4.1 | 3.1 | 55 | 4 | CALCIPRESSIN 1 |
| 1sb6-A | 4.1 | 2.4 | 50 | 8 | COPPER CHAPERONE SCATX1 |
| 3s1e-A | 4.1 | 2.4 | 52 | 6 | CYTOKININ DEHYDROGENASE 1 |
| 2g9o-A | 4.1 | 2.3 | 50 | 12 | COPPER-TRANSPORTING ATPASE 1 |
| 3v2u-C | 4.1 | 2.8 | 54 | 11 | GALACTOSE/LACTOSE METABOLISM REGULATORY PROTEIN G |
| 7t71-A | 4.0 | 4.3 | 53 | 6 | MEVALONATE 3.5-BISPHOSPHATE DECARBOXYLASE |
| 8csp-8 | 4.0 | 2.9 | 53 | 6 | 28S RIBOSOMAL PROTEIN S34. MITOCHONDRIAL |
| 6fon-A | 4.0 | 2.8 | 51 | 8 | COPPER CHAPERONE FOR SUPEROXIDE DISMUTASE |
| 6jsh-B | 4.0 | 2.9 | 55 | 11 | FATTY ACID SYNTHASE SUBUNIT BETA |
| 2l48-A | 4.0 | 3.2 | 57 | 5 | N-ACETYLMURAMOYL-L-ALANINE AMIDASE |
| 1ec6-A | 4.0 | 2.2 | 51 | 12 | 20-MER RNA HAIRPIN |
| 7np8-B | 4.0 | 3.5 | 54 | 15 | COENZYME F420-DEPENDENT SULFITE REDUCTASE |
| 4u9r-A | 4.0 | 3.4 | 55 | 16 | CZCP CATION EFFLUX P1-ATPASE |
| 4zck-A | 4.0 | 2.6 | 53 | 17 | GTP-BINDING PROTEIN TYPA/BIPA |

| | | | | | |
|---|---|---|---|---|---|
| 2wbm-A | 4.0 | 3.0 | 54 | 17 | RIBOSOME MATURATION PROTEIN SDO1 HOMOLOG |
| 3v97-A | 3.9 | 2.4 | 48 | 8 | RIBOSOMAL RNA LARGE SUBUNIT METHYLTRANSFERASE L |
| 7uvp-A | 3.9 | 3.4 | 55 | 9 | TETRACYCLINE RESISTANCE PROTEIN TETQ |
| 8hcn-D | 3.9 | 2.9 | 55 | 9 | UREASE SUBUNIT GAMMA |
| 2k1r-B | 3.9 | 2.6 | 49 | 6 | COPPER-TRANSPORTING ATPASE 1 |
| 3mah-A | 3.9 | 3.3 | 53 | 13 | ASPARTOKINASE |
| 8gh6-A | 3.9 | 3.4 | 53 | 8 | REVERSE TRANSCRIPTASE-LIKE PROTEIN |
| 6eml-p | 3.9 | 3.5 | 57 | 9 | PRE-18S RIBOSOMAL RNA |
| 7nad-U | 3.9 | 2.5 | 50 | 8 | 25S RRNA |
| 3oq2-B | 3.8 | 2.6 | 51 | 6 | CRISPR-ASSOCIATED PROTEIN CAS2 |
| 7agv-F | 3.8 | 3.5 | 52 | 2 | K(+)/H(+) ANTIPORTER SUBUNIT KHTT |
| 6cc2-A | 3.8 | 3.4 | 54 | 7 | CELL DIVISION CONTROL PROTEIN 45 CDC45 PUTATIVE |
| 5wwx-A | 3.8 | 2.9 | 52 | 10 | RNA-BINDING E3 UBIQUITIN-PROTEIN LIGASE MEX3C |
| 2ywg-A | 3.8 | 3.7 | 57 | 14 | GTP-BINDING PROTEIN LEPA |
| 5www-A | 3.8 | 2.5 | 54 | 11 | RNA-BINDING E3 UBIQUITIN-PROTEIN LIGASE MEX3C |
| 6yaq-A | 3.8 | 1.9 | 49 | 4 | CYTOKININ DEHYDROGENASE 8 |
| 5cwa-A | 3.8 | 4.1 | 58 | 7 | ANTHRANILATE SYNTHASE COMPONENT 1 |
| 3kiz-B | 3.8 | 2.5 | 52 | 8 | PHOSPHORIBOSYLFORMYLGLYCINAMIDINE CYCLO-LIGASE |
| 1eqr-A | 3.8 | 3.4 | 55 | 5 | ASPARTYL-TRNA SYNTHETASE |
| 5gan-G | 3.7 | 3.0 | 55 | 7 | SACCHAROMYCES CEREVISIAE STRAIN UOA_M2 CHROMOSOME |
| 4ct8-A | 3.7 | 3.6 | 57 | 14 | CINA-LIKE PROTEIN |
| 8d8j-F | 3.7 | 2.5 | 51 | 14 | PROBABLE S-ADENOSYL-L-METHIONINE-DEPENDENT RNA |
| 7ock-L | 3.7 | 2.1 | 51 | 6 | S-ADENOSYLMETHIONINE SYNTHASE |
| 7qri-A | 3.7 | 2.9 | 56 | 9 | TRYPTOPHAN 5-HYDROXYLASE 2 |
| 1ahu-A | 3.7 | 3.0 | 55 | 5 | VANILLYL-ALCOHOL OXIDASE |
| 1q5y-D | 3.6 | 2.7 | 52 | 8 | NICKEL RESPONSIVE REGULATOR |
| 6eld-A | 3.6 | 3.0 | 52 | 6 | NUCLEOLYSIN TIA-1 ISOFORM P40.U1 SMALL NUCLEAR |
| 6vej-A | 3.6 | 3.5 | 53 | 8 | PROBABLE RESISTANCE-NODULATION-CELL DIVISION (RND |
| 5yys-A | 3.6 | 3.3 | 54 | 7 | L-FUCOKINASE. L-FUCOSE-1-P GUANYLYLTRANSFERASE |
| 4yut-A | 3.6 | 3.3 | 54 | 6 | FAMILY 3 ADENYLATE CYCLASE |
| 6l5d-B | 3.6 | 3.1 | 52 | 6 | GAS VESICLE PROTEIN |
| 2pff-B | 3.6 | 2.9 | 54 | 11 | FATTY ACID SYNTHASE SUBUNIT ALPHA |
| 6fht-B | 3.5 | 3.8 | 55 | 5 | BACTERIOPHYTOCHROME.ADENYLATE CYCLASE |
| 6zvp-A | 3.5 | 2.3 | 52 | 4 | TYROSINE 3-MONOOXYGENASE |
| 1f93-A | 3.5 | 2.2 | 53 | 8 | DIMERIZATION COFACTOR OF HEPATOCYTE NUCLEAR |
| 7nhr-A | 3.5 | 3.2 | 53 | 15 | PUTATIVE TRANSMEMBRANE PROTEIN WZC |
| 6gwj-B | 3.5 | 2.6 | 52 | 8 | EKC/KEOPS COMPLEX SUBUNIT LAGE3 |
| 1v8c-A | 3.5 | 2.9 | 50 | 14 | MOAD RELATED PROTEIN |
| 1q8l-A | 3.5 | 2.5 | 52 | 8 | COPPER-TRANSPORTING ATPASE 1 |
| 3ui3-B | 3.4 | 2.3 | 50 | 8 | IMMUNOGLOBULIN G-BINDING PROTEIN G. VIRULENCE-ASS |
| 3c6k-D | 3.4 | 2.2 | 50 | 8 | SPERMINE SYNTHASE |
| 4qmf-B | 3.4 | 3.5 | 55 | 5 | KRR1 SMALL SUBUNIT PROCESSOME COMPONENT |
| 6nx5-A | 3.4 | 3.6 | 51 | 6 | PUMILIO DOMAIN-CONTAINING PROTEIN C56F2.08C |
| 6bwo-A | 3.4 | 2.8 | 49 | 8 | PYRIDINIUM-3.5-BISTHIOCARBOXYLIC ACID MONONUCLEOT |
| 4pwu-C | 3.4 | 3.0 | 53 | 8 | MODULATOR PROTEIN MZRA |
| 7qh2-C | 3.4 | 3.2 | 55 | 7 | LACTATE DEHYDROGENASE (NAD(+).FERREDOXIN) SUBUNIT |
| 6j6g-C | 3.4 | 3.4 | 55 | 7 | PRE-MRNA-SPLICING FACTOR 8 |
| 6s6b-K | 3.4 | 2.4 | 52 | 2 | CRISPR-ASSOCIATED PROTEIN. CMR5 FAMILY |

| | | | | | |
|---|---|---|---|---|---|
| 3afg-B | 3.4 | 3.2 | 56 | 13 | SUBTILISIN-LIKE SERINE PROTEASE |
| 4lir-B | 3.4 | 3.3 | 52 | 6 | NUCLEOPORIN NUP53 |
| 6teq-A | 3.4 | 3.0 | 55 | 7 | GALACTOKINASE |
| 2jvz-A | 3.4 | 2.4 | 52 | 8 | FAR UPSTREAM ELEMENT-BINDING PROTEIN 2 |
| 7m7h-B | 3.4 | 3.4 | 52 | 10 | ERYAI.6-DEOXYERYTHRONOLIDE-B SYNTHASE ERYA3. MODU |
| 4aim-A | 3.4 | 3.9 | 58 | 12 | POLYRIBONUCLEOTIDE NUCLEOTIDYLTRANSFERASE |
| 1u8s-B | 3.4 | 3.0 | 53 | 2 | GLYCINE CLEAVAGE SYSTEM TRANSCRIPTIONAL |
| 6pwn-A | 3.3 | 3.9 | 56 | 11 | SMALL-CONDUCTANCE MECHANOSENSITIVE CHANNEL |
| 3dkx-A | 3.3 | 3.3 | 55 | 15 | REPLICATION PROTEIN REPB |
| 7wvz-A | 3.3 | 3.5 | 51 | 10 | BETA-KETOACYL-ACYL-CARRIER-PROTEIN SYNTHASE I |
| 1fd8-A | 3.3 | 2.7 | 48 | 8 | ATX1 COPPER CHAPERONE |
| 2ko1-A | 3.3 | 2.8 | 53 | 8 | GTP PYROPHOSPHOKINASE |
| 4zos-A | 3.3 | 3.1 | 54 | 2 | PROTEIN YE0340 FROM YERSINIA ENTEROCOLITICA SUBSP |
| 6mrj-B | 3.3 | 2.8 | 53 | 11 | NICKEL-RESPONSIVE REGULATOR |
| 4olp-B | 3.3 | 2.4 | 49 | 4 | GRPU MICROCOMPARTMENT SHELL PROTEIN |
| 2gx8-C | 3.3 | 2.4 | 53 | 4 | NIF3-RELATED PROTEIN |
| 7v99-A | 3.3 | 3.3 | 53 | 4 | TELOMERASE REVERSE TRANSCRIPTASE |
| 6dgd-A | 3.3 | 2.7 | 53 | 8 | PRIMOSOMAL PROTEIN N' |
| 3wx4-A | 3.3 | 3.5 | 54 | 9 | ANTI-RESTRICTION ENDONUCLEASE |
| 1yqh-A | 3.3 | 2.5 | 52 | 2 | IG HYPOTHETICAL 16092 |
| 1vr6-A | 3.3 | 3.0 | 50 | 4 | PHOSPHO-2-DEHYDRO-3-DEOXYHEPTONATE ALDOLASE |
| 3gnw-B | 3.3 | 3.4 | 51 | 6 | RNA-DIRECTED RNA POLYMERASE |
| 1zav-A | 3.2 | 2.0 | 50 | 8 | 50S RIBOSOMAL PROTEIN L10 |
| 2hfs-A | 3.2 | 3.8 | 53 | 9 | MEVALONATE KINASE. PUTATIVE |
| 4zoq-F | 3.2 | 2.6 | 47 | 13 | INTRACELLULAR SERINE PROTEASE |
| 7qpr-D | 3.2 | 3.3 | 49 | 12 | ACT DOMAIN PROTEIN |
| 6s2e-A | 3.2 | 3.7 | 56 | 11 | DNA POLYMERASE EPSILON CATALYTIC SUBUNIT A |
| 2wbr-A | 3.2 | 3.6 | 53 | 8 | GW182 |
| 6pwj-A | 3.2 | 2.6 | 54 | 11 | GGDEF AND EAL DOMAIN-CONTAINING PROTEIN |
| 7jtk-i | 3.2 | 2.8 | 48 | 6 | FLAGELLAR RADIAL SPOKE PROTEIN 1 |
| 7cv0-A | 3.2 | 3.8 | 50 | 6 | TRANSCRIPTIONAL REGULATOR NIAR |
| 2f3j-A | 3.2 | 3.8 | 50 | 6 | RNA AND EXPORT FACTOR BINDING PROTEIN 2 |
| 5u9m-D | 3.2 | 2.4 | 46 | 9 | SUPEROXIDE DISMUTASE [CU-ZN] |
| 7q4l-A | 3.2 | 5.0 | 55 | 7 | DEAD END PROTEIN HOMOLOG 1 |
| 1kn6-A | 3.2 | 2.9 | 52 | 6 | PROHORMONE CONVERTASE 1 |
| 2hh2-A | 3.1 | 2.5 | 48 | 4 | KH-TYPE SPLICING REGULATORY PROTEIN |
| 5hb7-A | 3.1 | 3.2 | 53 | 8 | NUCLEOPORIN NUP53 |
| 7nhr-C | 3.1 | 3.5 | 54 | 15 | PUTATIVE TRANSMEMBRANE PROTEIN WZC |
| 4gzk-A | 3.1 | 2.7 | 50 | 12 | RNA-DEPENDENT RNA POLYMERASE P2 |
| 6wb2-A | 3.1 | 3.3 | 52 | 13 | HIV-1 VIRAL RNA GENOME FRAGMENT |
| 3opk-C | 3.1 | 3.3 | 56 | 4 | DIVALENT-CATION TOLERANCE PROTEIN CUTA |
| 6lpn-B | 3.1 | 3.6 | 54 | 13 | D-2-HYDROXYGLUTARATE DEHYDROGENASE. MITOCHONDRIAL |
| 7m1n-A | 3.1 | 2.9 | 47 | 9 | PUTATIVE FERREDOXIN |
| 3tvi-D | 3.1 | 4.0 | 55 | 15 | ASPARTOKINASE |
| 1qfr-A | 3.1 | 2.6 | 51 | 8 | PHOSPHOCARRIER PROTEIN HPR |
| 5uyy-A | 3.1 | 4.5 | 54 | 15 | PREPHENATE DEHYDROGENASE |
| 4usj-C | 3.1 | 3.1 | 53 | 0 | ACETYLGLUTAMATE KINASE. CHLOROPLASTIC |
| 8ba1-A | 3.1 | 3.7 | 54 | 6 | CLEAVAGE AND POLYADENYLATION SPECIFICITY FACTOR S |

| | | | | | |
|---|---|---|---|---|---|
| 8ily-A | 3.1 | 3.8 | 53 | 6 | SET DOMAIN CONTAINING 1A. HISTONE LYSINE METHYLTR |
| 3j6v-J | 3.1 | 3.0 | 55 | 7 | 28S RIBOSOMAL RNA. MITOCHONDIAL |
| 2raq-B | 3.1 | 2.8 | 53 | 9 | CONSERVED PROTEIN MTH889 |
| 3ihs-A | 3.1 | 3.3 | 52 | 8 | PHOSPHOCARRIER PROTEIN HPR |
| 1fx2-A | 3.1 | 3.1 | 55 | 7 | RECEPTOR-TYPE ADENYLATE CYCLASE GRESAG 4.1 |
| 7bbb-A | 3.1 | 3.3 | 50 | 8 | ATP-DEPENDENT RNA HELICASE DBPA |
| 4wd9-A | 3.1 | 2.9 | 53 | 11 | NISIN BIOSYNTHESIS PROTEIN NISB |
| 2ril-A | 3.1 | 2.6 | 49 | 12 | ANTIBIOTIC BIOSYNTHESIS MONOOXYGENASE |
| 6me0-C | 3.1 | 4.2 | 51 | 14 | T.EL4H RNA |
| 2lvw-A | 3.1 | 3.8 | 55 | 5 | ACETOLACTATE SYNTHASE ISOZYME 1 SMALL SUBUNIT |
| 1yg0-A | 3.1 | 2.3 | 45 | 11 | COP ASSOCIATED PROTEIN |
| 6cng-A | 3.0 | 2.7 | 41 | 10 | FATTY ACID KINASE (FAK) B3 PROTEIN |
| 1siz-A | 3.0 | 2.2 | 44 | 9 | FERREDOXIN |
| 8a8k-A | 3.0 | 3.1 | 55 | 4 | PAP PHOSPHATASE FROM METHANOTHERMOCOCCUS |
| 6ner-E | 3.0 | 2.8 | 52 | 6 | BMC-H TANDEM FUSION PROTEIN |
| 4p52-A | 3.0 | 4.1 | 58 | 7 | HOMOSERINE KINASE |
| 3mcs-B | 3.0 | 3.0 | 52 | 4 | PUTATIVE MONOOXYGENASE |
| 5anb-K | 3.0 | 3.0 | 52 | 10 | 60S RIBOSOMAL PROTEIN L3 |
| 4dnr-A | 3.0 | 2.5 | 53 | 9 | CATION EFFLUX SYSTEM PROTEIN CUSB |
| 1xpp-D | 3.0 | 3.1 | 52 | 13 | DNA-DIRECTED RNA POLYMERASE SUBUNIT L |
| 3ced-A | 3.0 | 3.6 | 55 | 13 | METHIONINE IMPORT ATP-BINDING PROTEIN METN 2 |
| 6lxg-A | 3.0 | 2.6 | 47 | 2 | GTP PYROPHOSPHOKINASE |
| 1fjg-F | 3.0 | 2.5 | 50 | 14 | 16S RIBOSOMAL RNA |
| 6dd5-A | 3.0 | 3.7 | 53 | 6 | MMB-1 CAS6 FUSED TO MALTOSE BINDING PROTEIN.CRISP |
| 4v1a-k | 3.0 | 2.5 | 53 | 4 | MITORIBOSOMAL PROTEIN ML37. MRPL37 |
| 7r65-A | 3.0 | 3.4 | 53 | 6 | ADENYLATE/GUANYLATE CYCLASE |
| 6k2e-A | 3.0 | 2.6 | 50 | 4 | CRISPR/CAS2 PROTEIN |
| 3d45-A | 3.0 | 3.8 | 53 | 11 | POLY(A)-SPECIFIC RIBONUCLEASE PARN |
| 5hy3-A | 3.0 | 2.0 | 47 | 15 | MRNA ENDORIBONUCLEASE LSOA |
| 5tl4-A | 3.0 | 2.9 | 53 | 4 | VANILLATE/3-O-METHYLGALLATE O-DEMETHYLASE |
